# Unbiased pangenome graphs

**DOI:** 10.1101/2022.02.14.480413

**Authors:** Erik Garrison, Andrea Guarracino

## Abstract

**Motivation:** Pangenome variation graphs model the mutual alignment of collections of DNA sequences. A set of pairwise alignments implies a variation graph, but there are no scalable methods to generate such a graph from these alignments. Existing related approaches depend on a single reference, a specific ordering of genomes, or a *de Bruijn* model based on a fixed *k*-mer length. A scalable, self-contained method to build pangenome graphs without such limitations would be a key step in pangenome construction and manipulation pipelines.

**Results:** We design the *seqwish* algorithm, which builds a variation graph from a set of sequences and alignments between them. We first transform the alignment set into an implicit interval tree. To build up the variation graph, we query this tree-based representation of the alignments to reduce transitive matches into single DNA segments in a sequence graph. By recording the mapping from input sequence to output graph, we can trace the original paths through this graph, yielding a pangenome variation graph. We present an implementation that operates in external memory, using disk-backed data structures and lock-free parallel methods to drive the core graph induction step. We demonstrate that our method scales to very large graph induction problems by applying it to build pangenome graphs for several species.

**Availability:** *seqwish* is published as free software under the MIT open source license. Source code and documentation are available at https://github.com/ekg/seqwish. *seqwish* can be installed via Bioconda https://bioconda.github.io/recipes/seqwish/README.html or GNU Guix https://github.com/ekg/guix-genomics/blob/master/seqwish.scm.

## 1 Introduction

A pangenome models the full genomic information of a species or clade (Medini *et al*., 2005; Sherman and Salzberg, 2020). In contrast to reference-based approaches that relate sequences to a particular reference genome, methods that use pangenome reference systems attempt to model the mutual relationship between all represented genomes (Consortium, 2018). Many approaches model the pangenome alignment as a *pangenome graph* (Garrison *et al*., 2018; Yokoyama *et al*., 2019; Hickey *et al*., 2020). A pangenome graph encodes DNA sequences as walks through an underlying language encoded in a sequence graph (Hein, 1989). In a pangenome graph, variation can be understood in the context of any part of any included genome (Eizenga *et al*., 2020). This lets us avoid the problem of reference bias, which can be understood as the limitation of analyses to genome sequences that are similar to a chosen reference genome.

An unbiased pangenome graph would represent the alignment of all included genomes to all others. Existing methods approximate this relationship by progressive alignment to a graph initially based on a reference genome (Li *et al*., 2020), through a global structuring of the genome relationships in a neighbor joining phylogenetic tree (Armstrong *et al*., 2020), or via creation of a *de Bruijn* graph based on a fixed *k*-mer length (Minkin *et al*., 2016; Sheikhizadeh *et al*., 2016; Yu *et al*., 2021). These methods limit computational costs by reducing the number of pairwise comparisons, but in turn their results depend on input genome order, selected reference, guide-tree topology, or *k*-mer length.

We consider the problem of building a pangenome graph without these potential sources of bias. Such a graph would be an ideal system to represent variation between two or more high-quality genomes. Given the rapid development of complete genome assemblies for humans and other vertebrates (Rhie *et al*., 2021; Nurk *et al*., 2021), we need a practical approach that can achieve this for tens to thousands of genomes on commodity hardware. Here, we present *seqwish*, an algorithm for the generation of a pangenome graph from pairwise alignments. Our solution is simple, but experiments on diverse sequence collections demonstrate that it easily scales to large pangenome building problems.

## 2 Algorithm

In this section, we provide a formal definition of variation graph induction. We then examine the bounds of a naïve implementation of this algorithm. Finally, we propose compression and partitioning techniques to reduce the space and working memory complexity of the induction process by a large constant factor modulated by the degree of sequence divergence in the input pangenome. This yields a practical algorithm for variation graph induction that can scale to the largest available pangenomes.

### 2.1 Variation graph induction

#### Definition 2.1.

Variation graphs are a common formalism to encode pangenome graphs (Garrison, 2019). In the variation graph 𝒢 = (𝒱, ℰ, 𝒫), nodes (or vertices) 𝒱 = *v*_1_ … *v*_|𝒱|_ contain sequences of DNA. Each node *v*_*i*_ has a unique identifier *i* and an implicit reverse complement 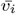. A node strand *s* corresponds to one node orientation. Edges ℰ = *e*_1_ … *e*_|ℰ|_ connect ordered pairs of node strands (*e*_*i*_ = (*s*_*a*_, *s*_*b*_)), encoding the base topology of the graph. Paths 𝒫 = *p*_1_ … *p*_|𝒫|_ describe walks over node strands 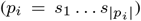, representing the collection of genomes embedded in the graph.

#### Theorem 2.1.

A variation graph represents pairwise alignments between its embedded paths.

Proof. By definition 2.1, two paths have identical subsequences where they walk (or step) through the same series of oriented nodes (e.g. *s*_1_*s*_2_*s*_3_). An identical set of path steps is thus equivalent to a sequence match. Pairwise alignments are by definition collections of character-level matches between sequences. The variation graph thus models a set of pairwise alignments between paths in 𝒫.

#### Theorem 2.2.

We can build a variation graph from sequences and pairwise alignments. The resulting variation graph fully embeds both the sequences and all pairwise relationships in the input.

This follows from 2.1. Our input 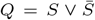 is a set of *N* DNA sequences *S* = *g*_1_ … *g*_*N*_ and their reverse complements 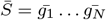. A match *m* = (*i, j*) asserts the aligned equivalence of two characters in sequences in *Q*. Pairwise alignments between sequences in *Q* are a set of matches *A* = {*m*_1_… *m*_|*A*|_}. By standard definition, each sequence matches its own reverse complement, that is 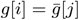 for all *j* = |*g*|−*i*, and we assume these matches are included in *A*. The transitive closure of a match, *m*^+^ = {*i* … *j*}, is a set of characters in *Q* that are transitively linked together by other matches. By definition of *m*, each *m*^+^ implies a single, identical character *c*(*m*^+^).

We build a graph 𝒢 inductively. We take the first match in *A, m*_1_, and execute a union-find operation to obtain 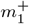. We add the character of the match 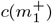 as a node *v*_1_ in 𝒢, and record the mapping from 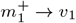. To induce the graph, we take the next unused match in 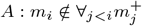, otain 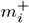, and add 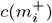 to 𝒢. To allow the annotation of paths, we record the set of characters in *Q* that match to a given node in 𝒢 in mapping *Z* = *Q* → 𝒱 = *m*_1_ … *m*_|𝒱|_. We continue until all matches have been used. Finally, we establish paths (𝒫) by walking them in 𝒢 using *Z*, and record edges (ℰ) where nodes occur successively in paths.

Proof. After the first step of induction, the graph represents all pairwise matches in 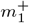. Each subsequent step includes progressively more of *A*, until at completion, all pairwise relationships are accounted for in *V*.

The set of alignments represented by a variation graph may be strictly larger than the set of alignments used to induce it. The graph must contain at least the set of alignments given in input. It may also contain new implied pairwise relationships that arise due to transitive match relationships, as shown in Figure 1 for closures 1 and 6. However, by definition of 𝒢, it cannot contain *less* match information than represented in the set of matches (*A*).

**Fig. 1:**
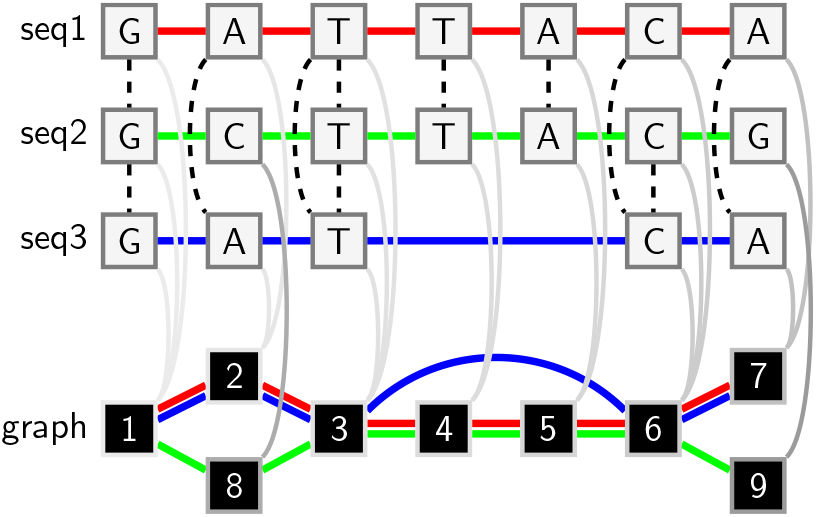
A visual description of variation graph induction. *Top*: an alignment graph model for three sequences and their alignments. Nodes are single characters (DNA base pairs, of which the forward strand is shown) in individual sequences. Solid edges link successive characters in each sequence, and are colored (red, green, blue) to identify each. Dashed edges indicate aligned pairs of characters. *Bottom*: a variation graph model induced from the alignment graph. Each transitive match closure (gray shaded edges, of increasing darkness) in the alignment graph results in a single node in the output graph, which is labeled by the rank of the transitive closure operation that produced it. By recording the full set of match closures, we can project the sequences in the input through to paths in the variation graph (colored edges). The unique set of node pairings in the paths provide the edges of the output graph. Closures 1 and 6 imply pairwise relationships between sequences seq1 and seq3, and seq1 and seq2, respectively, that are both absent in the input alignments.

### 2.2 Induction algorithm sketch

For the sake of time and space complexity analysis, we consider a simple algorithm to implement the induction process. The induction depends on our ability to compute transitive closures of matches *m*^+^. If *A* is sorted, we can find the matches of a given character in *Q* using binary search, which allows us to compute *m*^+^ for each character. We do so non-redundantly by marking each used character in *Q* in an auxiliary data structure 𝒳, which could be encoded as a bitvector of length |*Q*|. As we compute the transitive closures, we emit both the nodes (single characters) of the graph 𝒱 and the sequence-to-graph mapping *Z*, which, like *A*, consists of match pairs, but rather than mapping *Q* → *Q*, maps *Q* → 𝒱. As with *A*, we can sort *Z* to obtain random access via binary search. Finally, we derive elements in 𝒫 by iterating through the characters of *Q* and looking up their mapping in *Z* using binary search. The edge set ℰ are the unique pairs of steps found in 𝒫, and can be computed by sorting pairs of steps in 𝒫.

### 2.3 Naïve algorithm bounds

The inductive proof of theorem 2.2 demonstrates how to build a variation graph from sequences and their pairwise alignments. However, a naïve algorithm based on this model would require a very large amount of space. Although our identifier space |*S*| must include all of *Q*, in practice, we only store *S*, as 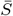 can be trivially computed. Assume an all-to-all alignment of *N* sequences in *A* as input, and that all sequences are approximately identical, so that the induced variation graph has |*S*|*/N* nodes. The induction must maintain reference to all characters in all input sequences *O*(|*S*|), all character-to-character matches *O*(|*A*|) ≈ *O*(|*S*|*N* ^2^), the mapping of *Q* into the graph *O*(|*Z*|) ≈ *O*(|*S*|), the nodes of the graph *O*(|𝒱|) ≈ *O*(|*S*|*/N*), the size of the edge set *O*(|ℰ|) ≈ *O*(|*S*|), and the set of paths *O*(|𝒫|) ≈ *O*(|*S*|). We also maintain the bitvector 𝒳 to mark seen characters of *Q* during graph induction, which requires *O*(|*S*|) bits, equivalent to *O*(|*S*|*/* log_2_ |*S*|) integer identifiers. In total, naïve theorem 2.2-based induction would require approximately *O*(|*S*|(*N* ^2^ + 1*/N* + 1*/log*_2_*Q* + 4)) space, or simply *O*(|*S*|*N* ^2^).

Assuming that we want to build a graph of 100 haploid human genomes of 3 × 10^9^ bp, where *N* = 100 and |*S*| ≈ 10^11^, we might expect to use ≈ 10^15^ identifiers to store the full model. Such a design is almost infeasible for inputs larger than a handful of genomes. For instance, we would need ≈ 3 ×10^12^ identifiers for just 5 human genomes. Although it is feasible to compute such a graph using external memory, the approximately 200-fold increase in space relative to the input renders this clearly impractical.

Considering the time complexity of induction, we anticipate *O*(|*A*| log |*A*|) time to sort the match set and *O*(𝒱 log |*A*|) to query it and compute our 𝒱 transitive closures. Computing the variation graph paths 𝒫 involves converting sequences in *S* to walks through 𝒱. We first sort the sequence-to-graph mapping array *Z* in *O*(|*S*| log |*S*|) operations, and then compute 𝒫 in *O*(|*S*|) queries which each cost *O*(log |*S*|). To obtain unique edges and generate ℰ, we must build and sort an array of size *O*(2|*P* |), and then iterate through it for *O*(2|*P* | log 2|*P* | + 2|*P* |) operations. In sum, we would expect to require *O*(|*A*| log |*A*| + V log |*A*| + 2(|*S*| log |*S*|) + 2(|*P* | log 2|*P* | + |*P* |)). Using our approximate relationships to |*S*| given previously and simplifying, we arrive at *O*[|*S*|(2*N* ^2^ log *N/* log |*S*| + (1*/N* + 4) log |*S*| + 2 + log(4))].

Due to our dependence on sorting, and the logarithmic-time cost of queries, growth in |*S*| drives Ω(|*S*| log |*S*|) growth in overall complexity. As *N* grows, both time and space complexity are dominated by the number of alignments, which in the case of our example is *O*(*N* ^2^). For large numbers of highly-similar genomes, we may not require all pairs of alignments to build a graph that contains all pairwise alignments. Various approaches could be used to reduce the size of *A* without disrupting the induced graph. We leave these to later work.

### 2.4 Match compression

As the bounds analysis shows, space requirements make it impractical to apply a trivial version of theorem 2.2 to generate a large pangenome graph. Therefore, we need a compression approach that exploits redundancy in the input genomes to reduce the costs of the algorithm. When working with large numbers of genomes, alignments dominate the computational costs. A simple technique is to generalize matches *m* = (*i, j*), which are between individual characters, to *range*-matches over pairs of ranges of characters in *Q*. For highly-similar sequences, our expectation is that exact matches will occur in long runs. If the average pairwise diversity of sequences in our input is 1*/k*, we expect exact matches to be around *k* characters long. By encoding matches as pairs of ranges of characters, *r* = (*a, b*) : *a, b* = (*i, j*) : *i, j* ∈ *Q*, we can obtain a ≈ *k*-fold compression of *A*, yielding the range-match array 𝒜.

If sorted, 𝒜 can be treated as an *implicit interval tree* (Li and Rong, 2020), which allows queries of containment and overlap in *O*(log |𝒜|) time. This compression requires trivial changes to our graph induction model. To obtain our match transitive closures (*m*^+^), we query 𝒜 for the range of a single character in *Q*, computing the character-level transitive relationships from the relative offsets of the ranges in 𝒜. Match compression thus reduces our alignment storage memory bounds by a factor of *k* without affecting our time complexity bounds.

The same encoding can be used to replace the sequence-to-graph mapping *Z*, yielding 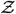. Rather than pairs of characters in *Q* and 𝒱, we record runs of matches between them as range matches. Although in expectation the length of these matches should be strictly less than *k*, due to the interruption of the graph by variation between genomes, this still allows us to reduce the size of *Z* using runs of matches between *Q* and 𝒱. Additionally, we store the inverse of 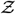, which maps ranges from 𝒱 → *Q*, As 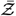. We use 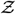 to compact non-branching regions of 𝒢 into single nodes, and 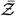 to accelerate our calculation of links in the graph.

### 2.5 Node compaction

For simplicity, we have thus far presented a character-level model of variation graph induction. However, range (or run) compression can also reduce the representation size of the graph. Rather than recording an identifier for each character in a sequence graph, it is useful to compact characters that form trivial linear components in the graph into single nodes. Broadly, the size of nodes will be bounded by the average distance between variants, which, for pangenomes built from ∼100 individuals of the same species, often provides a great reduction in the total number of nodes (and thus identifiers) required for 𝒢 and its components.

To compact 𝒢, we traverse *Q*, finding each entry in 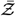 in turn, recording its start and end in 𝒱, which can be understood as a character vector or string containing all the sequence in the nodes of 𝒢. We subsequently use these markings to subdivide 𝒱 into a compacted version 𝒱′ where compacted node boundaries are marked in a auxiliary bitvector ℬ : |ℬ| = |𝒱| such that the first character in each compacted node is marked by a 1 and other characters are marked 0. ℬ allows us to compute compacted node ids using efficient rank operations (Gog *et al*., 2014).

### 2.6 Induction partitioning

Although match compression provides an approximate factor *k* improvement in memory bounds for key data structures used in the induction, the approach we present in section 2.4 requires working memory in the order of the set of transitive match closures in the graph. A simple approach to reduce this bound is to divide the induction problem into smaller pieces. We do so by computing the graph induction for a collection of initial characters in *Q*. In each partition, we apply a lock-free parallel union-find algorithm to derive the match closures (Anderson and Woll, 1991), appending results to appropriate data structures. This partitioning can introduce boundary effects which change the contents of 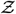 and 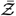 by splitting ranges at the boundaries of our partitions. However, while this will affect the compressed node definition 𝒱′, it does not affect V, and it can be corrected via a post-processing step to sort and compact the id space.

## 3 Implementation

We have presented a complete model for variation graph induction from sequences and their pairwise alignments (Algorithm 1). Here, we describe our specific implementation of this algorithm: *seqwish*. In general, our approach uses external memory to elaborate the graph, taking advantage of the availability of low-latency storage media, like solid-state drives (SSDs), to maximize the performance of this approach.

### 3.1 Input and output processing

Our implementation reads standard data formats, FASTA or FASTQ for the input sequences, and PAF (Li, 2018) for pairwise alignments. It writes the graph in standard Graphical Fragment Assembly (GFA) format (GFA Working Group, 2016).

#### Algorithm 1

The *seqwish* graph induction algorithm. For the sake of simplicity, we omit the details of several query algorithms that interact with the input alignments, the transitive match closure, implicit interval tree construction and query, node generation, and bitvector rank queries used in node compaction. Similarly, we omit the details of the input partitioning that we use to reduce maximum resident memory requirements.

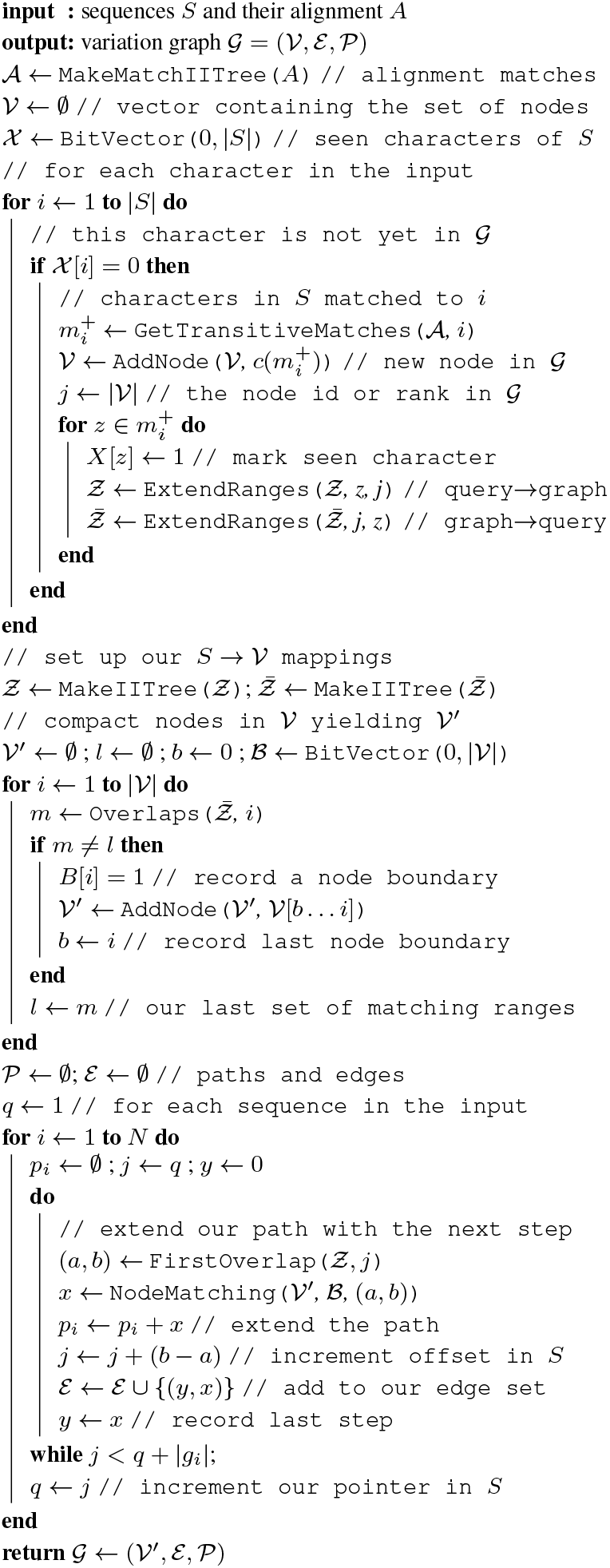

In PAF, the input set of alignments is not directly expressed in terms of matches between specific characters in *Q*. Rather, each record lists the name of the aligned pair of query and target sequences and offsets in each. To efficiently process the input PAF, we thus need to build a sequence index that allows us to generate A. In particular, we build a compressed suffix array (CSA) (Sadakane, 2000) over sequence names, that we call *seqidx*, and provide auxiliary supporting data structures that allow us to map between our input and the abstract concatenation of all input sequences and their reverse complements (*Q*). We often build graphs from very large collections of sequences, such as raw sequencing reads or contigs from many thousands of samples. This *seqidx* avoids the overheads associated with a hash table on string names of input sequences. To enable highly-efficient random access, we cache the input sequences in a disk-backed version of *Q*, into which our queries of sequence name and offset point. This trades time that might be spent accessing a compressed representation of the input for space in external memory.

For output in GFA, we iterate over nodes in 𝒱′, writing each as a node record. Edges are similarly produced from the disk-backed multiset representing ℰ. The most computationally expensive part of graph emission is the rendering of the input sequences *S* as paths 𝒫 through the graph. For each input sequence in the *seqidx*, we walk through the offsets in *S* contained in the sequence and look up their mapping into 𝒱′ using 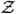. Range compression allows us to complete one lookup per range. By definition, each character in *S* is covered by only one range in 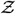. We can thus iterate through the ranges in 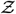 without considering each character. Following the GFA format, we are able to independently generate 𝒫, as each path is represented on a separate record in the GFA.

### 3.2 Key disk-backed data structures

In our implementation, we rely on several basic external memory kernels. To reduce working memory requirements to an absolute minimum, we use a disk-backed version of the implicit interval tree that memory-maps the sorted array of intervals (Garrison, 2021). Indexing the implicit interval tree requires a sorting step which dominates the runtime of our algorithm. We adapt the current best-performing in-place parallel sorting algorithm, In-place Parallel Super Scalar Samplesort (IPS^4^o), to work on a disk-backed, memory-mapped array (Axtmann *et al*., 2017). This allows us work with 𝒜, 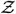, and 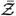 in external memory. By storing pairs of numerical identifiers in the backing array, we are able to generate a disk-backed multiset model which we use to compute the unique set of edges ℰ in terms of offsets in 𝒱. The graph sequence vector 𝒱 is simply written by appending characters to a file. We mark nodes to generate 𝒱′ using a bitvector kept in main memory, over which we subsequently generate a rank/select dictionary (Gog *et al*., 2014) for support of the final emission of the graph 𝒢.

### 3.3 Short match filter

Building a graph from an all-to-all alignment does not guarantee that the local structure of the graph is easy to understand. The all-to-all alignment is not coordinated, with each mapping aligned in isolation, and in consequence it fails to resolve the indel alignment normalization problem (Mose *et al*., 2019). This ambiguity can introduce deeply looping structures in the graph which collapse polymorphic microsatellites and other short VNTRs into very small numbers of nodes with very complex local topologies. Such motifs can cause problems with downstream analysis. We find that ambiguity about the arrangement of very short matches tends to drive complex local structures in the graph.

We mitigate this issue with a simple filter, *seqwish -k*, which simply ignores exact matches that are shorter than *k* characters. This filter necessarily increases the size of the induced graph. But, it also replaces complex motifs shorter than *k* with single bubbles. In doing so, it also removes short, expensive matches, reducing the overall space requirements for *seqwish*. When set very high, this filter can be used to generate a coarse, high-confidence graph built only from very long exact matches which will tend to be unique in the genome. Although the application of the *k >* filter can result in a graph that is relatively “under-aligned”, we can further refine it through the application of local multiple sequence alignment (Gao *et al*., 2020), or graph normalization (Doerr, 2022). In a pangenomic context, underalignment caused by *k >* match filtering can be mitigated by transitive relationships present in the pangenome.

## 4 Results

We evaluate *seqwish* through application to four pangenomes collected from *A. thaliana, H. sapiens, H. pylori*, and *Z. mays*. This limited survey is intended to demonstrate basic scaling properties of the method, and its practicality when applied to real pangenomes. We also consider the effect of the minimum match length filter described in section 3.3. Experiments were conducted on compute nodes with 386GB of RAM and AMD EPYC 7402P processors with 48 vCPUs.

To construct the graph we first generate alignments with *wfmash* (Guarracino *et al*., 2021), a DNA sequence aligner designed specifically for high performance all-to-all alignment of fully-assembled genomes. *wfmash* combines an algorithm for generating whole-genome homology maps (*MashMap2*) (Jain *et al*., 2018) with an extension of the wavefront algorithm (WFA) (Marco-Sola *et al*., 2020) capable of obtaining base-level alignments for whole chromosomes. *MashMap2* allows the user to define a homology length and pairwise divergence, expressed as a percent identity, over which to generate homology maps. This is useful when constructing pangenome graphs, because, in contrast to methods that are based on *k*-mer chaining (Harris, 2007; Li, 2018), it allows us to query the homology space of input genomes using two easily-interpretable parameters. The version of *wfmash* used in these experiments allows us to align sequences with up to 10% divergence between them, providing highly-sensitive input for our experiments. Our use of *wfmash* is pragmatic, and any mapping method capable of generating PAF with base-level alignments in CIGAR strings is capable of being used as input to *seqwish*.

In table 1 we provide input and constructed graph parameters for a single parameter setting of *wfmash* and *seqwish*, obtaining graph statistics with the ODGI toolkit (Guarracino *et al*., 2022). Figure 2 displays runtime versus graph size relative to the average input genome length across the range of parameters chosen for each pangenome. These provide a consistent set of insights. Reducing the sensitivity of alignments by increasing the identity threshold results in larger graphs. Filtering short matches results in larger graphs too, and for higher divergence collections of genomes, like *H. pylori*, tends to obliterate much of the homology information in the pangenome graph. This is visible from the fact that the “graph length / average genome length” ratio grows strongly as *k* increases. Such a reduction of the size of the set of matches considered for graph inductions also greatly reduces runtime. In all cases, we find that the initial alignment step takes longer than graph induction.

**Table 1:** Performance of the graph induction algorithm. For each pangenome we report a single experiment with *seqwish -k* filter set to 49bp. From left to right, the columns indicate the species, the number of sequences (that is, the number of contigs), number of haplotypes (that is, number of individuals), the sum of the length of all sequences in Gbp, the length of the short match filter applied in bp, the time in seconds and the amount of memory and disk space in Gbytes required for the graph induction, the length of the resulting graph in Gbp and the number its connected components.

| species | sequences | haplotypes | fasta.Gbp | min.match.bp | time.seconds | memory.Gbytes | disk.Gbytes | graph.Gbp | components |
| --- | --- | --- | --- | --- | --- | --- | --- | --- | --- |
| *A. thaliana* | 922 | 16 | 1.90251 | 49 | 468 | 43.1287 | 7.1218 | 0.234284 | 100 |
| *H. sapiens* | 17478 | 38 | 114.627 | 49 | 46268 | 347.4983 | 604.4261 | 4.47126 | 474 |
| *H. pylori* | 292 | 250 | 0.407782 | 49 | 777 | 74.9484 | 20.2070 | 0.01421 | 5 |
| *Z. mays* | 46289 | 41 | 90.2491 | 49 | 31043 | 351.1235 | 402.8716 | 13.8838 | 925 |

**Fig. 2:**
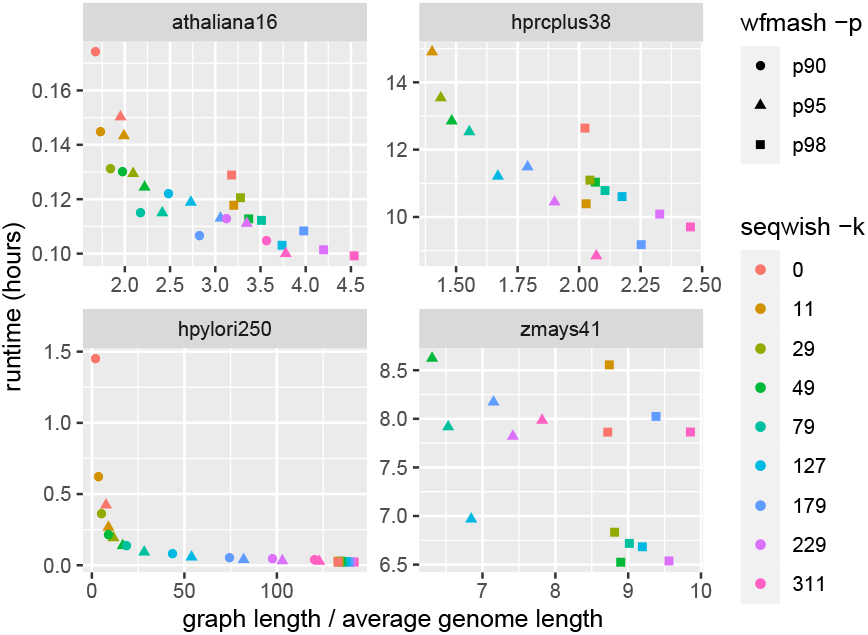
Experimental results from the application of *seqwish* to four different pangenomes. Each plot shows the runtime (in hours) versus the average input genome length calculated as the total length of all sequences in the pangenome divided by the number of included haplotypes. Multiple minimum identity settings for the mapping (*wfmash -p*) and different minimum match length filter settings (*seqwish -k*) result in a collection of graph builds per pangenome input. We compare the runtime in hours with the size of the resulting graphs relative to the average size of an input genome in the particular set. Lower limits on pairwise identity result in more compact graphs. Similarly, filtering short matches increases graph size relative to not (*seqwish -k*=0, red). The other way around, increasing the *seqwish -k* parameter tends to increase the size of the graph.

Although we use disk-backed data structures to represent the graph, the maximum memory requirements of *seqwish* are governed by the largest transitive match closure in the pangenome graph. We find that our particular partitioning scheme (we compact the graph in chunks as described in 2.6, using 50 Mbp chunks in all the experiments) does not allow us to complete the graph induction for *p* = 95 and *k* = 0 for the *H. sapiens* set, nor for *Z. mays* with *k* ≤ 29, where we run out of working memory. In practice, setting the chunk size lower tends to resolve this problem, but will also increase runtime. To simplify comparisons between the different parameter settings, we have not re-run these settings with a different partition size nor on computer nodes with a larger, then different, amount of RAM.

Our results with these four species-specific pangenomes give a basic demonstration of how pangenome graphs can be adjusted through key parameters of alignment and match filtering. These results are demonstrative of graph induction. In typical applications, we expect *seqwish* to be applied as part of a larger process to build and refine a pangenome graph (Garrison *et al*., 2022; Liao *et al*., 2022).

### 4.1 Comparison to related graph building methods

We have so far described our method for graph induction, provided detail on its implementation, and demonstrated its performance over a set of pangenome graph building problems. These provide intuition about *seqwish* and its behavior. However, the reader may wonder how the method compares to other similar methods for graph construction, or what variation might be caused by changes in the alignment method used.

To focus on these questions, we examine the case of a single chromosome (chrV) of *S. cerevisiae* for which we have 7 assemblies (Yue *et al*., 2017). We compute 100 random permutations (Williams, 2009) of these assemblies and provide them to several graph construction methods, including *minigraph* (Li *et al*., 2020), *TwoPaCo* (Minkin *et al*., 2016), and *seqwish* based on *minimap2* (Li, 2018) *or wfmash* (Guarracino *et al*., 2021) alignments.

Of existing methods, *TwoPaCo* provides a graph in a form that is similar to *seqwish*’s. However, it is based on a de Bruijn graph that must have a relatively low *k*-mer length, leading to complex topologies that in our experience cannot be resolved easily by increasing *k*. Furthermore, the overlaps between nodes are incompatible with the variation graph model, and must be reduced or “bluntified” with additional processing steps to be usable as input to variation graph tools (Eizenga *et al*., 2021). Because it is based on a de Bruijn graph, which is symmetrically constructed from a set of *k*-mers and not the sequences themselves, we do not expect *TwoPaCo* to be biased with respect to input genome order.

In contrast, *minigraph* develops a kind of partial order alignment (Lee *et al*., 2002) over the minimizers (sparse *k*-mer set) of input genomes, and it progressively builds the graph by adding new variation from each genome in order. We do expect it be order and reference (first input) biased, although the degree of bias requires experimentation to establish.

Finally, *seqwish* itself is by definition unbiased with respect to input order. But, if the alignments are affected by changes in order, the resulting graph will also change.

We use ODGI to compute the size, node count, and edge count of the resulting graphs (Guarracino *et al*., 2022). Changes in these statistics indicate changes in graphs. It also gives an indication of the scale of the differences, which may indicate how large an effect of a different construction approach might be. If graph parameters are the same for two large graphs, we consider them equivalent, which is not formally correct but provides a sufficient approximation for this analysis.

As we observe in Figure 3, *minigraph* indeed generates a distribution of graph lengths with clusters corresponding to permutations beginning with the same genome. This suggests that most of the variation in *minigraph*’s graph derives from the first genome which is picked. The graphs are thus biased towards the chosen base reference. However, this is not the case for the *seqwish* builds based on *minimap2* and *wfmash*, which appear to have almost exactly the same length across all permutations.

**Fig. 3:**
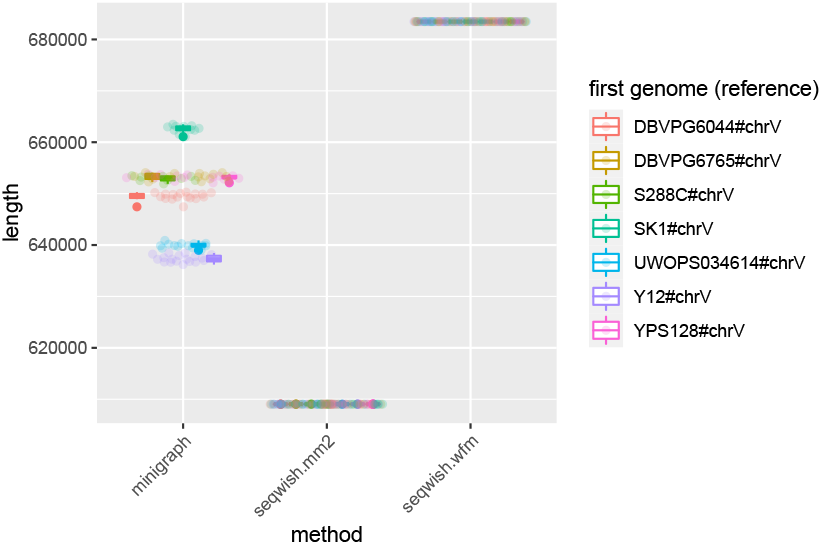
Base-pair lengths of graphs built from 100 permutations of yeast chrV. Each point corresponds to the length of a graph built from a specific permutation and the given method. We color each point by the first genome in the order (the “reference” in *minigraph*). A boxplot per first genome group provides the mean and the first and third quartile intervals for each reference grouping.

In Table 2, we further consider variance in the length, node count, and edge count of all graphs. This shows that, as expected, *minigraph*’s output depends markedly on input genome order. We also find that *minimap2* configuration we used is not order-independent, and can produce slightly different outputs when the reference and query sequences are ordered differently, demonstrating that *seqwish* is only “unbiased” insofar as its input mappings are. For *TwoPaCo* and *wfmash*+*seqwish*, we observe no variation in length. The length of *TwoPaCo* graphs is significantly larger due to the repetition in node sequences, which overlap by *k* − 1 base pairs.

**Table 2:** Variance in graph properties across genome input order permutations. We report the mean (*µ*) and variance in standard deviations (*σ*) for graph length (bp), node count (node), and edge count (edge). We show results from *minigraph, TwoPacCo* with *k* = 19 (*twopaco*.*k19*), *seqwish* using *minimap2* alignments (*seqwish*.*mm2*), and *seqwish* using *wfmash* alignments (*seqwish*.*wfm*).

| method | $\mu$.bp | $\sigma$.bp | $\mu$.node | $\sigma$.node | $\mu$.edge | $\sigma$.edge |
| --- | --- | --- | --- | --- | --- | --- |
| *minigraph* | 648967 | 8112 | 190 | 11.6 | 267 | 15.9 |
| *seqwish.mm2* | 609039 | 4.08 | 30692 | 2.89 | 41723 | 6.32 |
| *seqwish.wfm* | 683470 | 0 | 28756 | 0 | 39080 | 0 |
| *twopaco.k19* | 1270115 | 0 | 26383 | 0 | 35284 | 0 |

Although this particular study focuses on only a single small eukaryotic chromosome, it reveals that our basic understanding of reference bias in these graph construction methods is correct. By design, progressive pangenome graph construction methods will be affected by input genome order. But, other symmetric methods avoid order bias.

## 5 Discussion

We have presented a straightforward algorithm to generate a pangenome graph from a collection of genomes and alignments between them. By exploiting a simple model of this algorithm, we provide computational bounds that give insight into the complexity of the problem. We then make this approach practical by applying the concept of *match compression*, which reduces the expected computational complexity by a factor proportional to the diversity of input sequences. Our experimental results demonstrate that we can apply our method to various collections of sequences and alignments. It easily scales to some of the largest species pangenome construction problems possible using publicly-available, high-quality genome assemblies. *seqwish* is a generic sequence graph inducer of potentially many uses. We envision that it can serve as a component in diverse sequence analysis and assembly pipelines, and hope that our thorough description of its core algorithm and functionality will enable its reuse by other researchers.

It is also a potentially novel approach. Despite the existence of many methods for pangenome building, we are not aware of any comparable method which can losslessly convert an all-to-all alignment to a variation graph. This direct relationship allows users to adjust the shape of the resulting graph by modifying alignment parameters, allowing the design of custom graph construction processes based on domain-specific knowledge and potentially manual curation of assembly alignments. In contrast to existing methods, which depend on particular structuring of their input (Li *et al*., 2020; Armstrong *et al*., 2020), *seqwish* is unbiased in that as it directly and uniformly represents sequence relationships given on input in the resulting graph. Our comparison with related pangenome graph-building methods demonstrates that these effects can be significant. *minigraph*’s output varies substantially, with *σ* ≈ 1.25% of graph length for a 7-chromosome input with ≈ 99.3% pairwise identity. De Bruijn graph methods, which are also unbiased with their treatment of input genomes (Minkin *et al*., 2016; Sheikhizadeh *et al*., 2016; Yu *et al*., 2021).

Our presentation is necessarily limited, in order to focus on and describe the unique problem of variation graph induction. Thus, in this manuscript and our experiments, we have not explored the full problem of *pangenome graph building*, which include both the initial alignment step and downstream processing of the resulting graph (Liao *et al*., 2022). These topics lie outside of the scope of the presented work, wherein we have focused on a key kernel which is a bottleneck in the pangenome graph construction process. But, they are important for readers to consider. Although *seqwish* perfectly represents its input alignments, the problem of generating and filtering an alignment set remains critical, as it determines the structure of the built graph. And this lossless property does not guarantee that the resulting graph is easy to work with or navigate; in practice, downstream processing is usually required to normalize the graph for many applications. We expect to cover these topics in future work, some of which are ongoing (Liao *et al*., 2022; Garrison *et al*., 2022).

Despite its apparently high costs, the symmetric pangenome construction modality that *seqwish* presents provides key advantages over progressive approaches. Although a complete all-vs-all alignment of a set of genomes is costly, it is trivial to run in parallel. This feature is unavailable to progressive construction methods, which necessarily involve a serial introduction of new information into the graph. Although the total costs of a symmetric alignment and pangenome graph construction modality are high, they are in some sense more manageable. The modular, independent nature of the hardest part of the computation—the derivation of the alignments—can be scaled out to large compute clusters as well as cached for incremental construction and update of large pangenomes.

We furthermore expect that local giant components will tend to arise in the alignment graph without requiring a complete pairwise alignment, suggesting that lessons from graph theory may help to guide a justified sparsification, or downsampling, of the input (Janson *et al*., 1993). This suggests that simple random sampling of alignments may evoke minimal changes in graph structure, provided that there are many homologous copies of each locus in the pangenome. It will also be possible to apply *seqwish* to measure at what level of mapping sparsification we observe changes in the induced graph. Although this may be infeasible for very large problems, it will allow us to develop an empirical understanding of how to reduce the complexity of the initial alignment step without affecting the resulting pangenome graph.

## Acknowledgments

We are grateful to members of the HGSVC and HPRC production teams for their development of resources used in our exposition. We thank the authors of the pangenome resources made available on GenBank which have made our experiments possible.

## Data availability

Code and links to data resources used to build this manuscript and its figures can be found in the paper’s public repository: https://github.com/pangenome/seqwish-paper.

